# Resorbable alginate embolic microsphere for musculoskeletal applications

**DOI:** 10.1101/2025.01.13.632707

**Authors:** Sankalp Agarwal, Masum Pandey, Ronan Duggan, Con O’ Brien, Andrew L. Lewis, Brendan Duffy, Joan McCabe, Rena Daly, Liam Farrissey

## Abstract

Resorbable embolic agents can be used for musculoskeletal (MSK) pain relief applications using a Transarterial Embolisation (TAE) technique. However, inconsistent particle size and shape, degradation rates, and unpredictable resorption and excretion of the breakdown products, limit their widespread clinical adoption. The present study reports self-degradable, rehydratable, freeze-dried, resorbable Ca^2+^-crosslinked alginate microspheres as a novel vascular embolic agent. The alginate lyase enzyme was incorporated into the alginate microspheres to achieve a controlled degradation rate under physiological conditions. Whilst the feasibility of alginate microspheres as embolic agents has been demonstrated, this is the first study where the controlled release is achieved in physiological conditions. To incorporate the alginate lyase enzyme in the alginate matrix, its activity was reversibly inhibited by preparing an alginate lyase-alginate-excipient mixture in a pH 3.6–3.9 buffered solution. An electrostatic encapsulator enabled the scalable preparation of microspheres of size 210 ± 8 µm at room temperature. The results showed that freeze-dried microspheres regained their shape and size within two minutes after reconstitution with saline, along with active alginate lyase. The reperfusion time of alginate microspheres was evaluated using digital subtraction angiography (DSA) and compared with imipenem and celestine (IMP-CS)in a porcine renal arterial embolisation model. It was observed that most of the blood vessels reperfused within 130–210 minutes, and complete vascular blush was recovered within 24 hours. Embolization with resorbable alginate microspheres induced recoverable ischemic necrosis in the targeted renal arteries. This unique feature of resorbable alginate microspheres in inducing ischemic necrosis, potentially in pain-propagating neovessels, makes it an ideal candidate embolic agent for treating MSK-associated pain using the TAE technique.

## 1 Introduction

Transarterial embolisation (TAE) is an established minimally invasive techniquused to tree atof chronic K pain^1^. The painful MSK condition is thought to occur due to persistent tissue inflammation caused by the invasion of hyperandrogenic neovascularity and neoinnervation within the osteoarthritic joint space^2^. The neo-hyper vasculature supplies oxygen and nutrients to aberrant endothelial cells. It allows pro-inflammatory cytokines to infiltrate the local microenvironment, causing the tissue to become innervated and more pain-sensitive. TAE can be performed using occluding agents composed of either resorbable or non-resorbable materials, which prevent the blood flow at the target site and create ischemia-hypoxia induced necrosis of neovessels in addition to a decrease in the number of unmyelinated sensory nerves^3,4^. This breaks the pathological loop of neovascularisation, innervation, and chronic pain, reducing pain perception ^1,4^.

In clinical practice, the exact duration of necrosis depends on various factors, including the targeted vascular structure, the specific embolic material employed, and the clinical context of the embolisation procedure. However, neovessels are generally susceptible to ischemic damage because of their differentiated characteristics from normal blood vessels, such as intercellular gaps and altered basement membranes^3^. Researchers and clinicians have exploited this characteristic feature of neovessel to develop a new class of embolic agents that only transiently occlude the blood vessels to reduce hypervascularity and, subsequently, MSK-associated pain. There are various advantages of resorbable embolic agents over permanent embolic agents for MSK pain treatment ^5^. Firstly, due to the transient nature of the embolic agent, fewer off-target adverse events are likely to occur; secondly, transient occlusion may be sufficient to cause unmyelinated sensory nerve ischemia to relieve pain without creating prolonged localized numbness; and lastly, the use of resorbable agents allows for repeated TAE procedures to be conducted.

Many resorbable embolic materials are being evaluated in clinical practice for MSK pain relief, such as gelatin-based materials (sponge and microspheres), starch microspheres, and off-label IMP/CS particulate antibiotics^4,5^. Both gelatin- and starch-based embolic agents have shortcomings^5^. The gelatin sponge (such as Gelfoam® from Pfizer) must be cut manually, resulting in the formation of a polydisperse, irregular-shaped particle slurry, causing variability in the levels of occlusion. To overcome this problem, several manufacturers have developed gelatin microspheres (Nexsphere™ from Next Biomedical Co. Ltd, Korea, and IPZA™ from Engain C. Ltd, Korea). However, the gelatine-based agents suffer from unpredictability in degradation, as this relies upon local enzymatic action, which varies depending upon the quantity of embolic used, the site of embolization, and the patient’s biological response^6^. Indeed, some reports suggest a high proportion of permanent occlusion with gelatin-based embolic agents due to their small size particles^7^. Degradable starch microspheres (DSMs, such as EmboCept® S from PharmaCept GmbH, Berlin) are calibrated 50 µm particles of hydrolyzed starch crosslinked and substituted with glycerol ether groups having a very short half-life (< 40 minutes) ^3,8^. These are commonly used in combination with the delivery of chemotherapeutic drugs to liver tumours. However, they also suffer from unpredictable degradation rates when used in low blood flow anatomical territories, as their degradation depends solely on serum amylase. Off-label use of resorbable IMP/CS antibiotic particulates is widely used in clinical practice for pain relief in osteoarthritis, tendinopathy, adhesive capsulitis, and back pain. The recanalization time for IMP/CS ranges from 1 to 4 hours in vivo, which is considered an ideal duration to reduce neohypervascularity for MSK indications ^4^. Because IMP/CS particles are formed from an antibiotic mixture, its adoption as a temporary embolic device in mainstream clinical practice is hampered by concerns over antibiotic resistance development, allergic reactions, and significant regulatory challenges ^9^. For these reasons, there is a pressing need for a biocompatible resorbable embolic agent with a calibrated size and reliable resorption time. For its target product profile (TPP), the most appropriate size of microspheres for this use is considered-300 µm, as smaller particles can travel more distally and may result in unwanted off-target embolization to the skin and muscle. A clinical trial is being carried out using permanent Embozene microspheres of a similar size to treat knee pain secondary to moderate to severe osteoarthritis ^10^. As for the appropriate recanalization duration for temporary embolic materials, in vivo IMP/CS degradation time of 1-4 hours has been shown to be very effective in clinical practice and represents an appropriate target degradation rate for developing new resorbable embolic microspheres.

Alginate is a biocompatible polysaccharide previously used to prepare Ca2+-crosslinked microspheres for TAE ^11^. Under in vivo conditions, the backbone of the alginate may be degraded slowly and uncontrollably through non-enzymatic hydrolysis of glycosidic bonds. One strategy to enable both an increase in, and level of control over the degradation rate, is to incorporate an alginate-degrading enzyme within the alginate matrix. Leslie et al have demonstrated the immobilization of alginate lyase into alginate microspheres to achieve the tunable degradation of alginate microspheres for the controlled release of rat adipose stem cells to improve bone regeneration ^12^. However, several technical challenges were not addressed in the reported work to enable its usage for novel applications such as TAE, methods to preserve the product and provide a shelf-life, and to ensure its sterility for safe administration into the human body. Here, we have therefore, focused on developing stable, reconstitutable, freeze-dried alginate lyase-loaded Ca^2+^-crosslinked alginate resorbable microspheres that can be terminally sterilized. This study addresses an important challenge, ensuring that alginate lyase-entrapped alginate microspheres can be produced using medical-grade alginate for a sufficient length of time to achieve a good yield and can be preserved at room temperature without being degraded by the enzyme. Other aspects related to using these microspheres as resorbable embolic agents are discussed in this study: (i) characterization of alginate microspheres for the degradation time in vitro, (ii) determining the degradation time of resorbable microspheres in vivo and (iii) evaluating its effectiveness for inducing recoverable necrosis in renal arteries. To the best of our knowledge, this is the first report demonstrating the development of the resorbable alginate lyase-loaded alginate microsphere as an embolic agent for MSK pain relief applications.

## 2 Material and methods

### 2.1 Preparation of lyophilized resorbable alginate beads

The method of preparing alginate microspheres using an electrostatic encapsulator was adopted from Leslie et al, with appropriate modifications ^12^. The alginate beads were prepared by dissolving 2% w/v low-viscosity LVM sodium alginate (Novamatrix Pronova®, Oslo, Norway) in 0.1 M pH 2.8 acetate buffer, mixed 0.15U alginate lyase (Nagase Cooperation, Japan) solution to achieve the desired degradation rate. The acetate buffer inhibits the alginate lyase degrading activity, thereby allowing the synthesis of beads without losing the viscosity of alginate over a 30-minute time period at room temperature. thus, improving the throughput of resorbable microspheres. The beads of size around 200 µm were prepared through electrostatic encapsulation. The microspheres were then washed with a proprietary mixture of cryoprotectants and transferred into lyophilizer for freeze drying (Biopharma Process Systems Ltd, Winchester, UK) at -45^°^C degrees Celsius for 90 hours ^13^. The lyophilized microspheres were sealed under a vacuum and sterilized.

### 2.2 In vitro degradation of resorbable alginate microspheres

This is a quantitative bespoke method that was developed to simulate the degradation of microspheres in vivo. The lyophilized microspheres made with 0.15U was reconstituted with 1.5 mL of saline for 2-3 minutes, followed by an addition 6 ml of 0.1M pH 7.4 HEPES buffer (supplemented with physiological levels of calcium, phosphate, and sodium ions). The hydrated beads were filtered using a 20µm pore filter and removed the excess liquid before weighing not more than 110 mg (100mg, +10/0) of beads as a starting quantity. The weighed beads were dispersed into 6 ml HEPES filtrate, incubated at 37 ^°^C and gently agitated at 150 rpm (to avoid any mechanical damage) to evaluate the mass loss. The residual mass was recorded every 20 minutes until it reached less than 10 mg of weight. Where necessary, the 6 mL test volume of HEPES was maintained throughout the test period. A volume of 6 mL of HEPES Buffer was selected as it is sufficient to facilitate repeatable rinsing through a filter and creates an environment with sufficient ion exchange to degrade the microspheres to the level required for this test.

### 2.3 Animal model and husbandry

The swine model was selected for this study. The healthy swine model was chosen as the experimental species for this study because the size and anatomy of the vascular system are clinically relevant for the purpose of testing interventional medical devices. The porcine model has been widely used for vascular and physiology experiments to evaluate MSK embolization. The study was reviewed and approved by an external ethical committee for compliance with regulations prior to study initiation. Animals were individually or group housed in conditions meeting or exceeding the requirements outlined in the Directive 2010/63/EU and as described in the Guide for the Care and Use of Laboratory Animals.

#### 2.3.1 Embolization of the Renal arteries

In this study, 5 Yorkshire swine weighing 55-60 kg were used. After induction of anesthesia, using a standard Seldinger’s technique, the femoral artery was punctured, and a 6 Fr arterial sheath (Merit Medical, USA) was placed. 100-200 IU/kg of heparin were administered intravenously before selective catheterization for the test article delivery. Under fluoroscopic guidance, a guide catheter (Medtronic Inc., USA) was advanced through the sheath over a guide wire (Abbott, USA) into the target arteries. A microcatheter (Medtronic Inc., USA) was coaxially inserted through the guide catheter. The catheter tip was positioned at the origin of the target site. A vial of sterile resorbable microspheres was reconstituted with 1.5 mL of saline and 2 minutes was allowed to dissolve the contents to obtain a clear solution of hydrated beads. To the hydrated beads, 2 mL of non-ionic contrast agent (Omnipaque, GE healthcare) was added and gently inverted several times to mix the solution. A sterile 21-gauge needle was inserted to withdraw the beads in a 1 mL sterile syringe. Then, the needle was removed, and the syringe was attached to the catheter to start with injections of beads. Across 5 animals, 10 injections were delivered with aliquots ranging from 0.8 mL to 1.4 mL of resorbable alginate microsphere solution to target sites in kidneys under angiographic guidance to achieve 100% complete stasis in all the injections.

A similar procedure was followed to evaluate the embolization of IMP/CS (Imipenem/cilastatin antibiotics) in the kidneys of three animals. A suspension of IMP/CS was prepared by adding 5 ml of non-ionic iodinated contrast in 0.5 g of IMP/CS, followed by 10 secs of vigorous mixing, and was injected with 0.2 mL increments into the 6 different regions of the kidneys across 3 animals to achieve complete stasis. Angiography was performed at various follow-up times to assess the reperfusion. Animals were sacrificed and kidneys were collected and sent for histopathology analyses.

### 2.4 Histology

Out of 5 animals embolized with resorbable microspheres, 3 animals were euthanized on day 1 and the remaining on day 3 by intravenous injection of a barbiturate-based euthanasia solution^11^. The animal on day 1 was sacrificed to determine the presence of the embolic material as well as their biological fate to correlate with the in vitro studies (SEC-MALS). The kidneys were prepared and immersed in 10% NBF for the histology processing. For IMP/CS, three animals were sacrificed on day 3 of the survival period. In the case of IMP/CS, the objective was not to identify the presence of the embolic material but to identify the possibility of observing tissue necrosis. The harvested kidneys were trimmed to document to assist in maintaining orientation. The tissues were divided into proximal, middle and distal treated regions. The trimmed sections were fixed and embedded in paraffin blocks, sectioned and stained with hematoxylin and eosin ^11^. Using light microscopy, histological evaluations were performed to identify the presence of microspheres or their fragments, haemorrhage, inflammation and revascularization.

### 2.5 Statistical analysis

All the experiments were performed at least in triplicate. The statistical significance was tested for p < 0.05 using On-way ANOVA with post hoc Tukey test. The values were calculated as mean ± S.D.

## 3 Results

### 3.1 Preparation and in vitro degradation of resorbable alginate microspheres

Using optimal conditions, an acceptable particle size of 210 ± 8 µm encapsulated with 0.0875U enzyme of size approximately 200 µm at a high throughput were prepared as shown in Figure 1.

**Figure 1.**
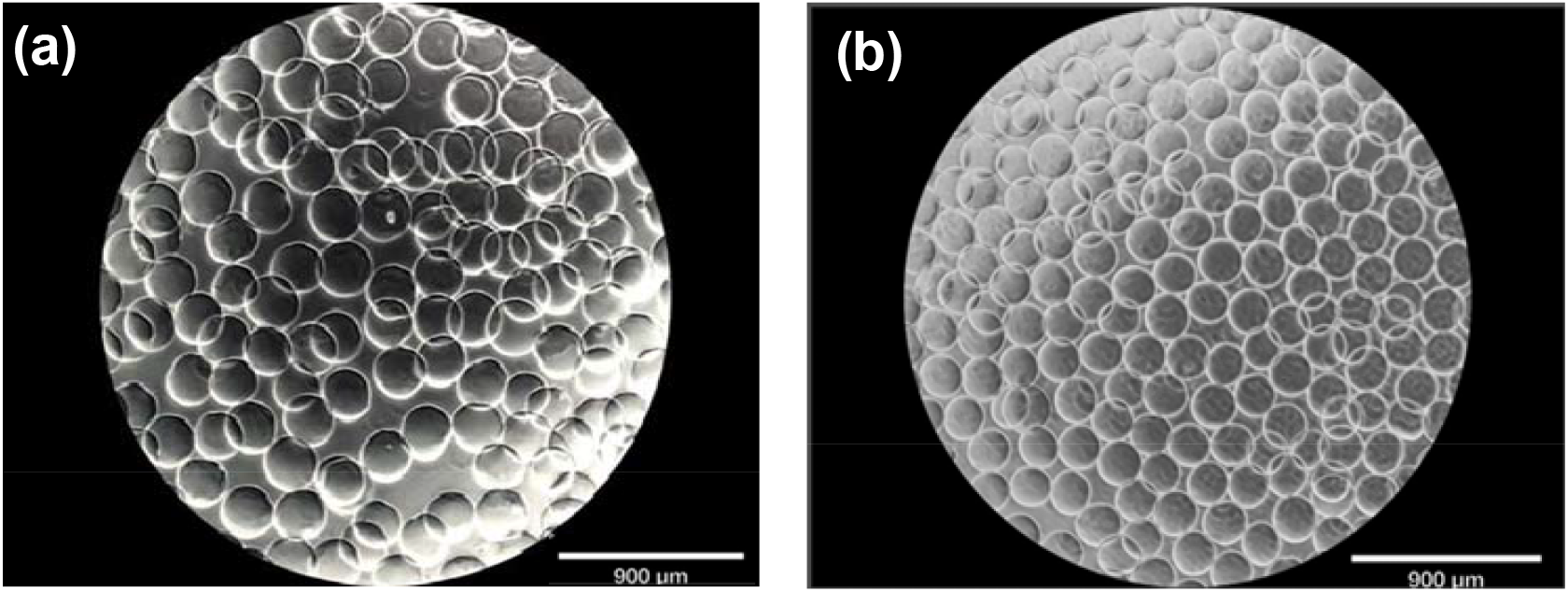
Representative images of pre-lyophilized (a, b) and post-lyophilized rehydrated/reconstituted microsphere encapsulated with 0.15 U alginate lyase enzyme, respectively.

The reconstituted microspheres made with 0.0875U alginate lyase were degraded in simulated in vitro conditions shown in Fig 2(a). The 10 mg value was selected as it is more difficult to accurately detect a change as the mass approaches zero due to the measurement accuracy of the equipment. The degradation time range for 0.0875U resorbable alginate microsphere is shown in Fig 2 (b) which is in the range of 140-240 minutes.

**Figure 2.**
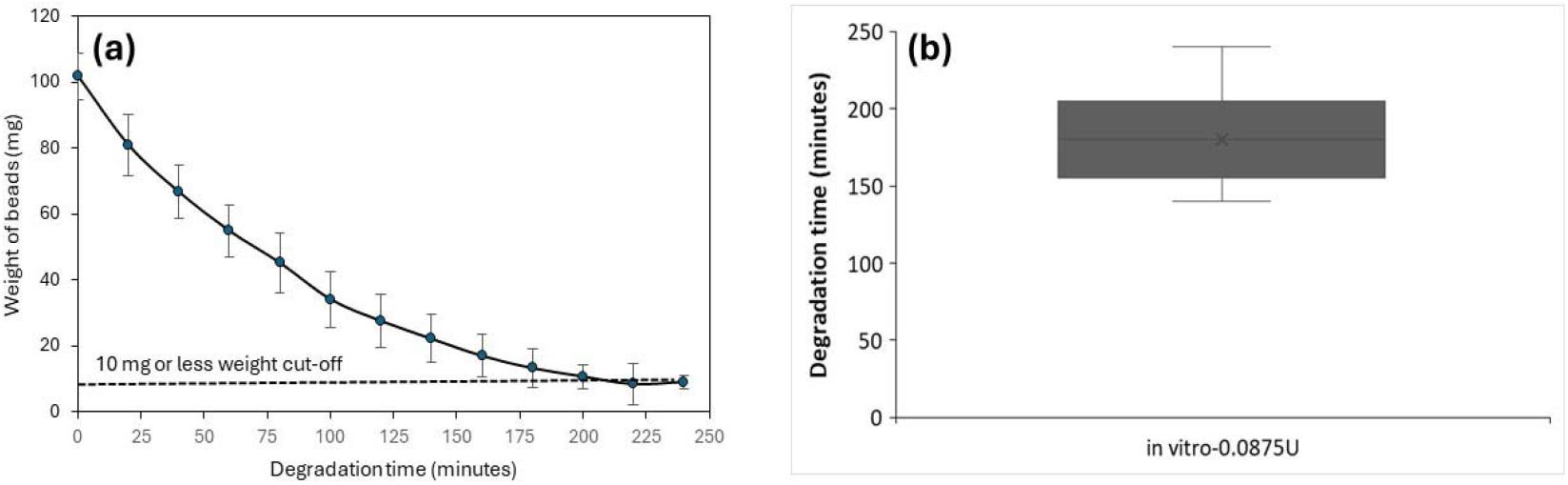
In vitro degradation of resorbable microspheres (a) Degradation profile to achieve volume reduction by 99.9% and (b) the degradation time.

### 3.2 Porcine renal arteries embolization

The representative DSA images comparing the embolization with resorbable alginate microspheres and IMP/CS, and their reperfusion times are given in Fig 3 (I and II). Here, the reperfusion time in vivo is defined as the time required for the complete reperfusion of the parent or main targeted arteries. The renal embolization sites for resorbable microspheres and IMP/CS are shown in Fig 3I(a) and 3II (a) respectively (denoted by yellow colour arrows). The main or parent arteries were reperfused at 150 minutes for both resorbable microspheres and IMP-CS (indicated by green arrows) whereas the grey blush area representing the more distal and smaller capillary network was partially reperfused (denoted by orange colour arrows). At 24 hours, complete revascularization of all the embolized vessels can be observed for resorbable microspheres and IMP/CS antibiotic particles. These results indicate the similarity of the IMP/CS particles and resorbable microspheres in occlusion reperfusion times. The range of reperfusion times of the resorbable microsphere was compared with IMP-CS as shown in Fig 3 (III). In addition, the in vivo reperfusion time of RM was compared to the in vitro degradation time. It has been observed that the reperfusion time of RM is in the range of 130-210 minutes and was found to be statistically similar to IMP-CS reperfusion time range of 60-247 minutes. Also, in vitro degradation time of resorbable microsphere was found to be statistically similar to in vivo degradation time. This demonstrates that the method employed to evaluate the in vitro degradation simulates the in vivo reperfusion trends/conditions, thus it can be used to predict the degradation of resorbable microspheres.

**Figure 3.**
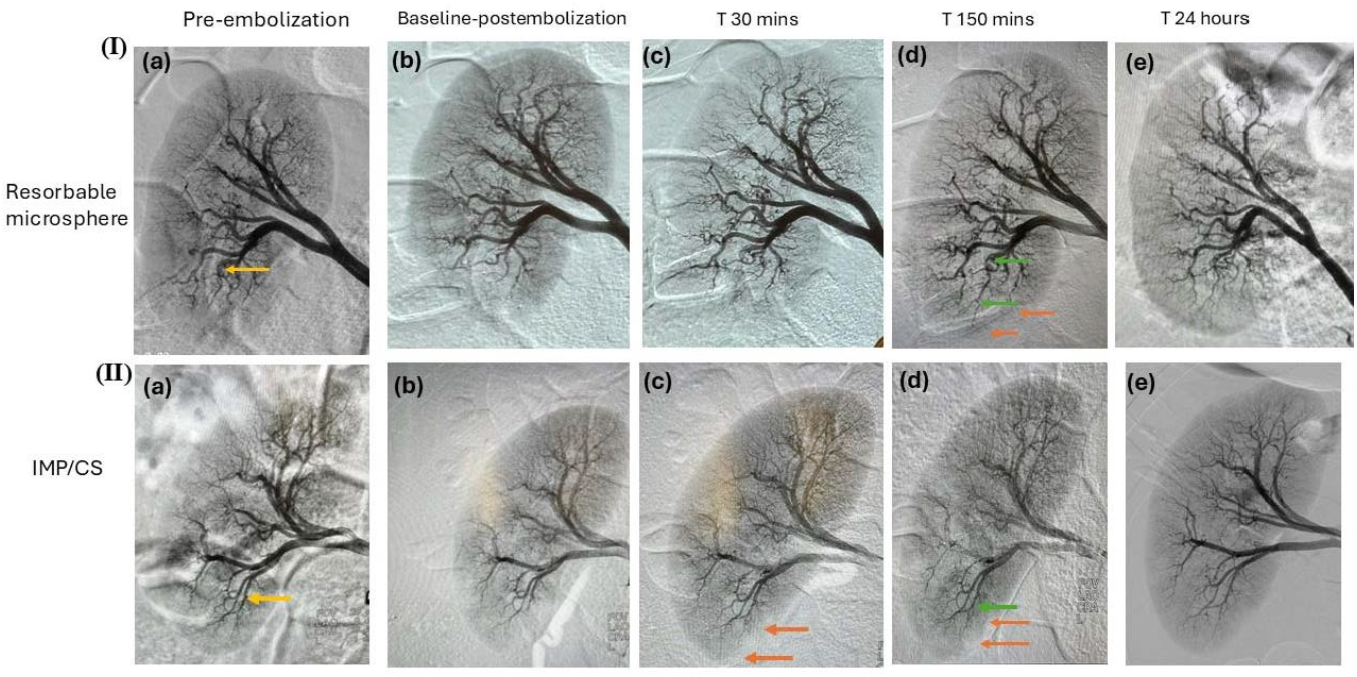
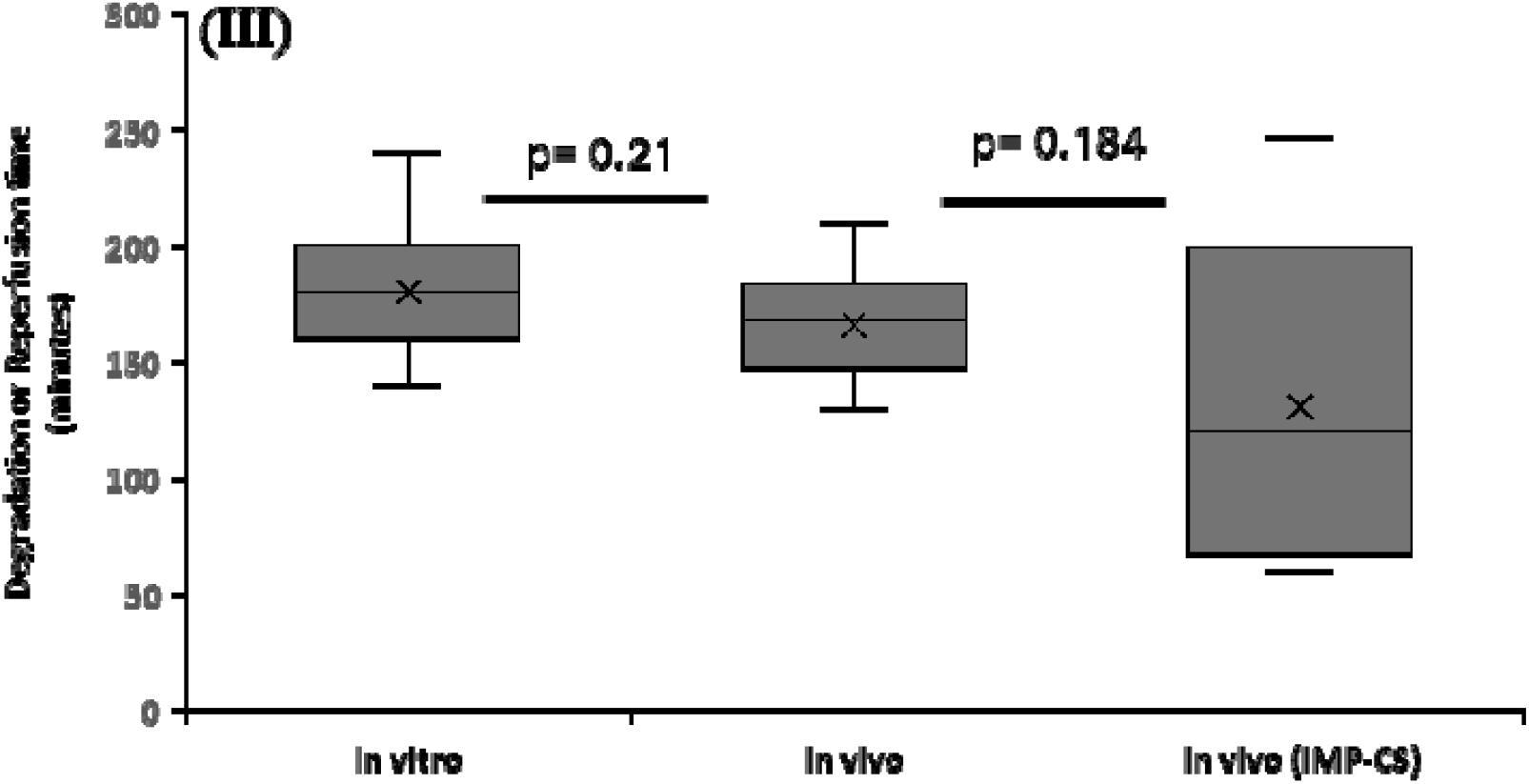
Representative of angiography images showing the occlusion and reperfusion of renal arteries in a porcine animal over 24 hours using (I) Resorbable alginate microsphere and (II) IMP-CS as embolic agents. (III) Comparison of in vivo reperfusion time and in vitro degradation time of resorbable microspheres (0.0875U) and IMP. (Site of embolization is denoted by yellow arrows, reperfusion of main or parent arteries is denoted by green arrows and grey blush is denoted by orange arrows).

Representative histology images of renal tissue embolized by resorbable alginate microspheres and IMP/CS are shown in Fig 4. No trace of embolic material was observed on day 3 histology as shown in Fig 4 (A). Likewise, the same observation can be made for the day 3 histology of IMP/CS treated kidney as shown in Fig 4 (B).

**Figure 4.**
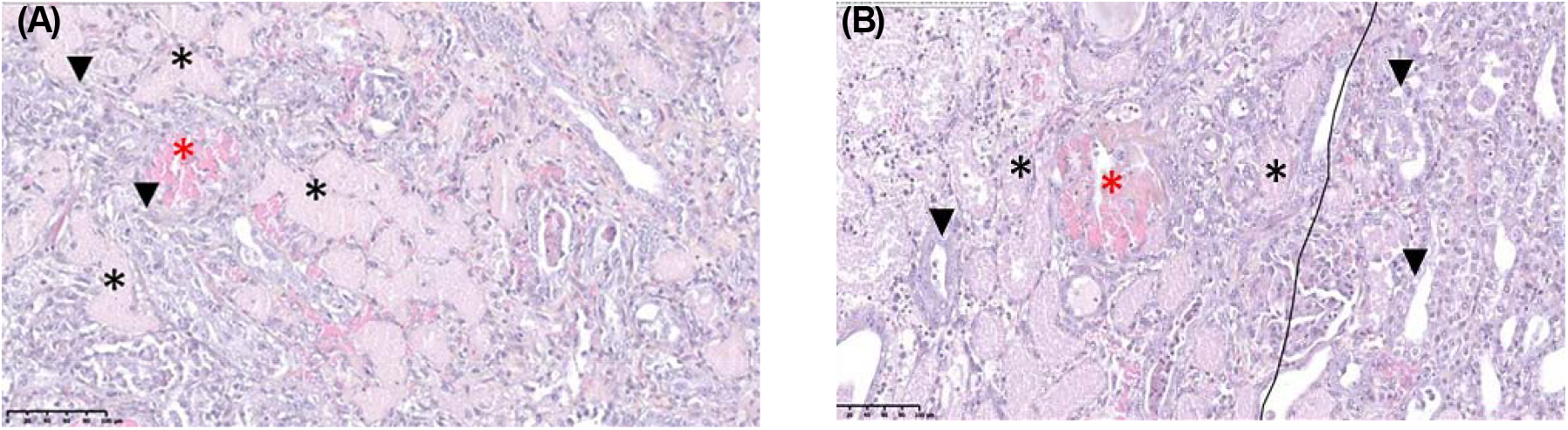
Representative histology images of renal tissue treated with resorbable microspheres after 72 hours (with a higher magnification image on the right) (A) and (B) IMP-CS. (Denotations: Renal tubular necrosis*, glomerular necrosis* and regeneration▾)

No trace of embolic material was observed on day 3 histology as shown in Fig 4 (A). Likewise, the same observation can be made for day 3 histology of IMP/CS treated kidney as shown in Fig 4 (B). The day 3 histology of resorbable microspheres shows ischemic lesions with renal tubular necrosis (stars), characterized by renal tubules with few epithelial cells and a homogeneous pink color and glomerulus necrosis (denoted by red stars). Close to these necrotic tubules, tubules under regeneration are observed (denoted by triangles). These tubules have more mitotic epithelial cells indicating the regeneration of renal tissues. The same observation can also be made from the day 3 histology of IMP-CS treated kidney.

## 4 Discussion

Resorbable alginate microspheres used for the transient embolization of blood vessels to create recoverable ischemic necrosis must be prepared through a process that ensures stability at the point of care for the clinical applications. Both the rate and extent of degradation should be controlled to ensure timely excretion of breakdown products from the body to ensure the ensemble is suitable for clinical application. Leslie et al previously demonstrated the preparation of alginate lyase-loaded alginate microspheres, however, several drawbacks of their system render their clinical use as an embolic agent untenable. For instance, no attempts were made to inactivate the enzyme to enable the long-term storage of the alginate microspheres until needed at the point of clinical care. Additionally, the feasibility of preparing microspheres over a prolonged time to facilitate adequate manufacturing throughput was not addressed. Kunjukunju et al demonstrated the loading of native or an aggregated form of alginate lyase in alginate microspheres^14^. They also adopted the use of a freeze -drying method (albeit without the use of any excipients or cryoprotectants) to preserve the microspheres for the release of biologics. In their study however, they failed to disclose if the microspheres fully recovered their shape and size after reconstitution; this is an important prerequisite for embolic microspheres, as they must flow through a narrow lumen microcatheter without clumping or blockage during delivery but must subsequently embolize blood vessels predictably and efficiently. The present work used a proprietary excipient mixture, alginate and alginate lyase dissolved in the acetate buffer of pH 2.8. This approach not only reversibly inhibits the alginate lyase activity during the synthesis of microspheres, enabling efficient enzyme encapsulation, but also provides stability to the alginate matrix during the lyophilization process to prevent the collapse of the structure. This process produces an elegant lyophilized cake containing resorbable microspheres which can be sterilized using gamma-radiation. The microspheres can be constituted using saline in less than 2 minutes, offering ease of use for clinicians that fit within their existing clinical workflow. Compared to the existing literature, the method of preparing alginate lyase-loaded calcium ion-crosslinked alginate microsphere is very unique and results in a product well-suited for clinical applications.

The IMP-CS antibiotic mixture has been widely used as an off-labelled embolic agent for MSK pain relief applications. The reported literature shows that an occlusion time of 1-4 hours is effective in inducing necrosis of neovessels ^4^. Therefore, the present work targeted a similar embolization time by investigating the incorporation of 0.0875U of alginate lyase in the alginate microspheres. The degradation time of 0.0875U microspheres is also more closely aligned to the upper end of the IMP-CS resorption time in vivo.

In the renal kidney embolization model, the embolization was induced by intra-arterial injection of resorbable alginate microsphere and IMP-CS as shown in Fig3I(b) and II (b) respectively. The parent arteries reperfused at the same time for both the embolic agents (Fig 3 I (c) and II (c)), around 150 minutes and a complete reperfusion the distal capillary blush was observed at 24 hours. The range of reperfusion times of resorbable microsphere and IMP-CS were comparable as shown in Fig 3 (III). The variation in the reperfusion times of the resorbable microsphere is affected largely by the anatomy of arteries and the depth of the embolization. The narrower blood vessels and distal embolization with resorbable alginate microspheres increase the reperfusion time due to the limited availability of fluid to drive the ion-exchange effect. In the case of IMP-CS, the variation in the reperfusion time is quite broad due to the polydisperse nature of IMP-CS particle. This makes it difficult to inject IMP-CS at the target site in a controlled and repetitive manner, leading to variation in the resorption rate of the particles in the blood. Furthermore, the in vivo reperfusion time of resorbable alginate microspheres was compared with the in vitro degradation time to determine the predictability of the microsphere degradation through in vitro method (Fig 3 III). The degradation time under in vivo and in vitro conditions for microspheres was shown to be statistically similar. Thus, it can be said that there is a good correlation between the in vitro degradation method and the in vivo conditions.

To determine the effect of embolization on the surrounding tissues, a gross histological evaluation of resorbable alginate and IMP-CS was performed on day 3 post embolization. On day 3, no resorbable alginate microsphere embolic material was observed in any sections, though necrotic and regenerative tissues were observed, as shown in Fig 4(a). In Fig 4(b), day 3 histology of IMP-CS treated kidneys showed histological observations similar to those of the resorbable alginate microspheres. Therefore, through angiography, it can be seen that the vessel gets revascularized completely at 24 hours occluded with resorbable microspheres, whereas recoverable necrosis with no presence of embolic material can be seen on day 3.

## 5 Conclusion

A resorbable calcium-crosslinked alginate lyase loaded-alginate microsphere was developed successfully as an embolic agent for MSK pain relief applications. A robust and scalable method was developed to encapsulate the enzyme and produce a gamma sterilized and rehydratable freeze-dried resorbable alginate microspheres. Controlled degradation was achieved under physiological conditions by incorporating alginate lyase into the alginate microspheres.

## 7 Acknowledgement

This study was co-funded by the European Union and Enterprise-Ireland DTIF grant. Views and opinions expressed are, however, those of the authors only and do not necessarily reflect those of the European Union or European Innovation Council and SMEs Executive Agency (EISMEA). Neither the European Union nor EISMEA can be held responsible for them.

## 8 Conflict of Interest statement

Liam Farrissey reports financial support was provided by Disruptive Technologies Innovation Fund (Grant number: 174147), Enterprises Ireland. Liam Farrissey reports a relationship with CrannMed.

## Notes

### Summary of Updates

The manuscript has been revised and updated

